# Interneuron dysfunction in a new knock-in mouse model of SCN1A GEFS+

**DOI:** 10.1101/849240

**Authors:** Antara Das, Bingyao Zhu, Yunyao Xie, Lisha Zeng, An T. Pham, Jonathan C. Neumann, Grant R. MacGregor, Soleil Schutte, Robert F. Hunt, Diane K. O’Dowd

**Author notes:** Department of Biophysics and Physiology, University of California, Irvine, CA 92697. Cold Spring Harbor Laboratory, Cold Spring Harbor, NY 11724. These authors contributed equally (Bingyao Zhu, Yunyao Xie).

## Abstract

Advances in genome sequencing have identified over 1300 mutations in the SCN1A sodium channel gene that result in genetic epilepsies. However, how individual mutations within SCN1A produce seizures remains elusive for most mutations. Previous work from our lab has shown that the K1270T (KT) mutation, which is linked to GEFS+ (Genetic Epilepsy with Febrile Seizure plus) in humans, causes reduced firing of GABAergic neurons in a *Drosophila* knock-in model. To examine the effect of this mutation in mammals, we introduced the equivalent KT mutation into the mouse *Scn1a* (*Scn1a*^*KT*^) gene using CRISPR/Cas9. Mouse lines carrying this mutation were examined in two widely used genetic backgrounds, C57BL/6NJ and 129×1/SvJ. In both backgrounds, homozygous mutants had spontaneous seizures and died by postnatal day 23. There was no difference in the lifespan of mice heterozygous for the mutation in either background when compared to wild-type littermates up to 6 months. Heterozygous mutants had heat-induced seizures at ~42 deg. Celsius, a temperature that did not induce seizures in wild-type littermates. In acute hippocampal slices, current-clamp recordings revealed a significant depolarized shift in action potential threshold and reduced action potential amplitude in parvalbumin-expressing inhibitory interneurons in *Scn1a*^*KT/+*^ mice. There was no change in the firing properties of excitatory CA1 pyramidal neurons. Our results indicate that *Scn1a*^*KT/+*^ mice develop seizures, and impaired action potential firing of inhibitory interneurons in *Scn1a*^*KT/+*^ mice may produce hyperexcitability in the hippocampus.

## Introduction

Mutations in *SCN1A*, which encodes the α subunit of the voltage-gated sodium channel Nav1.1, result in a broad spectrum of genetic epilepsies (Catterall, 2012; Miller and Sotero de Menezes, 2019; Schutte et al., 2016). Among 1300 known epilepsy causing *SCN1A* mutations, approximately 10% of the missense mutations are associated with genetic epilepsy with febrile seizures plus (GEFS+) (Miller and Sotero de Menezes, 2019; Zhang et al., 2017). Individuals with GEFS+ exhibit seizures induced by fever (febrile seizures) that begin in infancy but persist beyond six years of age (hence, febrile seizures plus or FS+). Some GEFS+ patients develop additional seizure phenotypes later in life including afebrile generalized tonic-clonic, myoclonic or absence seizures (Abou-Khalil et al., 2001; Escayg et al., 2001; Meng et al., 2015; Scheffer, 1997; Schutte et al., 2016; Zhang et al., 2017). The mechanism underlying the GEFS+ disorder associated with the different missense mutations remains poorly understood. Moreover, variations in seizure severity can occur even in family members with identical *SCN1A* mutations (Abou-Khalil et al., 2001; Zhang et al., 2017) suggesting that environmental factors and/or genetic modifiers (Bergren et al., 2009; Hawkins and Kearney, 2016; Hawkins et al., 2011; Jorge et al., 2011) can affect the phenotype. This has made developing effective therapies for treating GEFS+ patients challenging.

Animal models with specific missense mutations are important in evaluating the cellular and circuit mechanisms of GEFS+. Previously we generated a knock-in fly line, carrying the GEFS+ K1270T mutations, and identified alterations in sodium currents that reduce firing specifically in inhibitory neurons, contributing to a heat-induced seizure phenotype (Sun et al., 2012). Additionally, a rat model of GEFS+, with the N1417H mutation, has hyperthermia-induced seizures and reduced action potential amplitude in hippocampal interneurons (Hayashi et al., 2011; Mashimo et al., 2010). However, the only well-studied mammalian genetic knock-in model for SCN1A GEFS+ to date is the R1648H mouse line. In this GEFS+ model, animals exhibit heat-induced seizures as well a sleep, cognitive and social behavior deficits. The electrophysiological properties of neurons, role of genetic modifiers and therapeutic testing have been evaluated using this mouse model (Dutton et al., 2017; Hawkins et al., 2011; Hedrich et al., 2014; Martin et al., 2010; Papale et al., 2013; Salgueiro-Pereira et al., 2019; Sawyer et al., 2016; Wong et al., 2016). The key cellular mechanism findings posit that the mutant mice exhibit heat-induced seizures due to impaired firing in inhibitory neurons (Hedrich et al., 2014; Martin et al., 2010).

Given the heterogeneity of GEFS+ disease phenotypes, additional mouse models will be important in understanding the cellular mechanisms contributing to specific aspects of disease caused by different missense mutations. In this study, we used CRISPR/Cas9 to generate a mouse model of the GEFS+ SCN1A K1270T mutation that we previously studied in a knock-in *Drosophila* model. Mutant K1270T mice exhibit heat-induced seizures and a low incidence of spontaneous seizures, similar to human patients and mutant K1270T fly phenotypes (Abou-Khalil et al., 2001; Sun et al., 2012). Comparison of mutant mice in two genetic backgrounds (C57BL/6NJ vs 129×1/SvJ) revealed no strain differences in lifespan or heat-induced seizures. Further, electrophysiological analysis showed the K1270T mutation reduces the excitability of parvalbumin (PV)-expressing interneurons in the mouse hippocampus, while there were no significant differences in the excitability of pyramidal neurons. These findings are partially consistent with *Drosophila* and human induced pluripotent stem cell (hiPSC) models of K1270T generated by our lab (Schutte et al., 2016; Sun et al., 2012; Xie et al., 2019), as well as other *Scn1a* mouse models, that implicate reduced inhibition as a likely cause of hyper-excitable neural networks which in turn, gives rise to seizures (Cheah et al., 2012; Dutton et al., 2013; Hedrich et al., 2014; Kalume et al., 2013; Martin et al., 2010; Ogiwara et al., 2007; Rubinstein et al., 2015a; Sun et al., 2012; Tai et al., 2014; Xie et al., 2019; Yu et al., 2006). This new knock-in mouse model broadens our ability to explore how distinct missense mutations alter cellular and circuit mechanisms giving rise to SCN1A GEFS+ phenotypes.

## Materials and Methods

### 1. Animals

Mice were maintained on a 12-h light/dark cycle with *ad libitum* food (Envigo-Teklad global rodent diet #2020X) and water. All experiments were performed in accordance with the Institutional Animal Care and Use Committee of the University of California, Irvine.

*Scn1a* K1270T mutant mice were generated by CRISPR/Cas9 and maintained on a C57BL/6NJ (JAX Stock # 005304) or 129×1/SvJ (JAX Stock # 000691) background. A full description of the targeting of K1270T is described below. In some experiments, *Scn1a*^*KT/+*^ mice were mated with a PV-Cre (JAX Stock # 017320) and Ai14-tdTomato reporter (JAX Stock # 007908) on a C57BL/6J background to generate wild-type and heterozygous mice that have parvalbumin (or PV) neurons endogenously labeled with fluorescent red tdTomato marker. Experiments were performed on male and female littermates between ages 3-4 month for EEG studies and between P28-P40 for all other experiments. No sex differences were observed for lifespan or seizure behavior.

### 2. CRISPR/Cas9

Mice encoding *SCN1A* K1270T were generated by introducing a missense mutation within exon 19 at an equivalent position (K1259T) in mouse *Scn1a* via CRISPR/Cas9 in combination with a single stranded oligodeoxynucleotide (ssODN) for use in homology dependent repair (HDR). DNA template for synthesis of a single guide RNA 72 (sgRNA72) was produced by PCR using in order, forward primer (TMF372), T7 ribopromoter, 20bp guide sequence, region of homology to plasmid pSLQ1651-sgTelomere(F+E) (Addgene #51024; (Chen et al., 2013)). The reverse primer (TMF159) binds to the plasmid and defines the 3’ end of the single guide RNA (sg72). Sequence of the forward and reverse primers were: TMF372 (5’ – gttaatacgactcactatagTCCTGGAGATGCTCCTCAAAgtttaagagctatgctg-3’), and TMF159 (5’ – GAAAAAAAGCACCGACTCGGTGCC – 3’). After purification of the PCR product, T7 RNA polymerase was used to synthesize the sgRNA. Messenger RNA encoding Cas9 was synthesized using T7-mediated *in vitro* transcription of pT7_pX330 followed by capping and poly-adenylylation as directed by the manufacturer (mMessage mMachine T7 ultra transcription kit, Thermo-Fisher). B6SJLF2/J fertilized mouse oocytes were microinjected with sgRNA72 (10ng/ul), Cas9 mRNA (10ng/ul) and a modified single-stranded deoxyribonucleotide (ssODN) (Integrated DNA Technologies, Ultramer) as a template for HDR. The ssODN contained the desired sequence to introduce the K1259T amino acid change, silent base changes to reduce likelihood of re-cutting of the desired allele, plus a silent base change to generate an *EcoR*V site for screening. Surviving oocytes were transferred to the reproductive tract of pseudo-pregnant CD1 females. After screening of offsprings, a male with the desired modification (#1472) was used to establish lines of *Scn1a* K1259T mice on a C57BL/6NJ (JAX stock# 005304) or 129×1/SvJ (JAX Stock # 000691) strain background by backcrossing with wild-type females for at least five generations.

### 3. PCR

To screen for the *Scn1a* K1270T mutation, DNA extracted from toe biopsies (Lysis buffer and proteinase K, Viagen) was amplified using a *Scn1a*-specific forward primer (TMF 441-TTCCATCCCAAGGAAATACCATGT) and reverse primer (TMF442 - GCCTATCTTGTCATCACAACACAGTG). PCR amplification was performed for 35 cycles of 94°C for 30s, 59°C for 30s and 72°C for 30s using recombinant 5U/ul Taq polymerase prepared in 10X Buffer with 50mM MgCl_2_ (Invitrogen). The 388bp PCR amplicon was digested with 20U/µL EcoRV restriction enzyme prepared in 10X NEB 3.1 buffer (New England Biolabs) and incubated at 37^°^C for 60min to distinguish between the wild-type (388bp) and mutant (223bp and 165bp) alleles.

### 4. Heat-induced seizures

Wild-type and *Scn1a*^*KT/+*^ mice (P30-P40) of both sexes and weighing ≥15g were used to screen for occurrence of heat-induced behavioral seizures. Each mouse was briefly anesthetized using isoflurane and a rectal temperature probe (RET 3, Braintree Scientific) was stably inserted and secured to the tail. The probe was attached to a thermometer (TW2-107, Braintree Scientific) which was used to monitor core body temperature throughout the assay. A custom-built heating chamber was preheated to 50°C for at least 1 hour prior to the start of the experiment to ensure uniform heating of the chamber (Figure 3A). The chamber is equipped with a digital temperature controller to set the chamber to a desired temperature. A thermocouple fitted inside the chamber gives feedback to the temperature controller to stably maintain the chamber temperature to the set value. Each mouse, fitted with a rectal temperature probe, was introduced into the preheated chamber. The chamber temperature was increased by 5°C every 10 min (Figure 3A, bottom panel) using the digital temperature controller. Core body temperature of the mouse was recorded every minute throughout the experiment. Each mouse was video monitored and seizure severities were scored offline by an experimenter blind to mouse genotype, based on a modified Racine scale (Racine scoring scale 1-mouth and facial movements; 2-head nodding; 3-forelimb clonus, usually one limb; 4-forelimb or hindlimb clonus, hindlimb extension, falling on side; and 5-excessive jumping, generalized tonic-clonic seizure). For each animal, the seizure threshold temperature, latency to seizure and seizure behavior score was recorded. The experiment was terminated when 1) the mouse had a seizure, or 2) body temperature reached 44°C. At the end of the assay, the mouse was placed on a pre-chilled dish to rapidly restore normal body temperature. All experiments were done between 10:00 A.M and 4:00 P.M; the experimenters were blind to the genotype.

### 5. EEG monitoring

Wild-type and heterozygous and mice between 3-4 month-old were observed using EEG monitoring system (Pinnacle Technologies) as previously described (Kim et al., 2018). Briefly, each mouse was anesthetized with ketamine and xylazine (10 mg and 1 mg per kg, intraperitoneal) so that there was no limb-withdrawal response to a noxious foot pinch. Prefabricated EEG headsets (#8201, Pinnacle Technologies) were surgically implanted overlying the hippocampus, and cemented in place with a fast-acting adhesive and dental acrylic. Two wires were laid on the shoulder muscles for electromyographic recording. Mice were allowed to recover for 5-7 d before recordings were initiated. After recovery from surgery, each mouse was placed in a cylindrical plexiglass chamber for EEG recordings. Experimental mice were monitored for 7-9 d (24 h d^−1^). Electrographic EEG seizures episodes were manually detected by scanning the raw data files using Sirenia Seizure software (Pinnacle Technologies) while the experimenter was blind to mouse genotype. Electrographic seizures were defined as high-frequency, high-voltage synchronized polyspike waves with amplitudes at least twofold greater than background that lasted ≥15 s.

### 6. Immunostaining

Mouse brains were harvested from P30-P40 wild-type and heterozygous mice with genetically labeled PV neurons (*Scn1a*^*+/+*^*; PV-Cre; tdTomato* or *Scn1a*^*KT/+;*^ *PV-Cre; tdTomato*) and sectioned into 250-300 µm using vibratome (VT1200, Leica). Brain slices were fixed with 4% paraformaldehyde (PFA) for 15 min at room temperature (RT) followed by overnight incubation at 4 °C. After multiple washes with 1X PBS (phosphate buffer saline), brain slices were blocked in PBS-BTA (4% BSA+ 0.25% TritonX100+ 0.02% sodium azide prepared in 1XPBS) for 1 hr followed by incubation with primary antibody-anti-Parvalbumin (1:1000, P3088 Sigma-Aldrich) at RT for 4 hrs with gentle shaking, and overnight incubation at 4 °C. Excess antibody was removed with several washes with 1X PBS before incubation with secondary antibody anti-mouse conjugated with Alexa Fluor 488 (1:1000, A11029 Life Technologies) for 2-4 hrs at RT with gentle shaking. Excess antibody was removed by multiple 1X PBS washes. Brain slices were stained with DAPI (1:40000, Sigma-Aldrich) for 15 min at RT with gentle shaking followed by with several washes with 1X PBS. Brain slices were mounted with Fluoromount G (0100-01, Southern Biotech) and imaged with a LSM700 Zeiss confocal microscope using a 20X and 40X objectives. Brightness and contrast were adjusted, and images were assembled using Zeiss ZEN software and Adobe Photoshop.

### 7. Whole cell electrophysiology

Brains were quickly harvested from deeply anesthetized wild-type and *Scn1a*^*KT/+*^ C57BL/6NJ littermates (P28-40), and 300-350 µm coronal brain slices were prepared in oxygenated ice-cold high-sucrose ACSF (artificial cerebrospinal fluid) containing: 150mM sucrose, 124 mM NaCl, 3 mM KCl, 1.25 mM NaH_2_PO_4._H_2_O, 2 mM MgSO_4._7H_2_O, 26 mM NaHCO_3_, 10 mM dextrose and 2 mM CaCl_2_. Slices were kept at 32 °C for at least 30 min prior to electrophysiological recordings in Axopatch 700B, Digidata 1440A and pCLAMP10.1 or 10.2 software. Electrophysiological recordings were performed at room temperature (25 °C) with glass pipettes with open tip resistance of 3-6 MΩ (TW150F-3, World Precision Instruments, & 100µL calibrated pipettes #534332-921,VWR International). The bath solution consisted of ACSF: 124 mM NaCl, 3 mM KCl, 1.25 mM NaH_2_PO_4._H_2_O, 2 mM MgSO_4._7H_2_O, 26 mM NaHCO_3_, 10 mM dextrose and 2 mM CaCl_2_. The electrode was filled with internal solution either 1) 120 mM potassium gluconate, 20 mM NaCl, 0.1 mM CaCl_2_, 2 mM MgCl_2_, 1.1 mM EGTA; 10 mM HEPES, 4.5 mM Na_2_ATP (pH= 7.2), or 2) 140 mM potassium gluconate, 1 mM NaCl, 1 mM CaCl_2_, 1 mM MgCl_2_, 5 mM EGTA, 10 mM HEPES, 3 mM KOH and 2 mM ATP (pH =7.2). Osmolarity of external ACSF and internal solution was adjusted to 310-312 mOsm and 288-290 mOsm respectively using VAPRO 5200 osmometer (Wescor). Signals were obtained using pCLAMP10.2 with a Bessel filter at 2-4 KHz and a low-pass filter at 10KHz. The passive properties of the cell were determined in voltage clamp mode. Resting membrane potentials were measured immediately after break-in with no current injection at current clamp mode. To record firing properties, cells were injected with a series of long (1000ms) current injections starting at −80pA with 10pA step increment. Range of current injection steps was-80pA to 200pA for excitatory neurons and −80pA to 900pA for PV inhibitory neurons. The number of action potentials fired in 1000ms was compared between wild-type and heterozygous mice. Data analyses was performed using pCLAMP Clampfit10.6 and Microsoft Excel.

### 8. Quantification and Statistics

Survivorship curves were compared between wild-type, heterozygous and homozygous mice with log-rank Mantel-Cox test. Mouse mean body weights for each genotype was calculated with at least three mice per time point. Mean body weight among homozygous, heterozygous and wild-type mice was compared with Mann-Whitney test for each time point. Body weights between heterozygous and wild-type mice in respective strain background was compared using two-way ANOVA with genotype and postnatal age (days) as factors, followed by Sidak’s post hoc multiple comparisons (*p* < 0.05).

Rate of change of body temperature across time, seizure threshold temperature, seizure latency, Racine score etc. was compared between wild-type and heterozygous mice using two-tailed unpaired Student’s t-test or Mann-Whitney test as specified in the text. Incidence of seizure occurrence was compared between genotypes with Fisher’s exact test.

For each individual cell, first action potential fired was used to calculate threshold, amplitude and half-width using pCLAMP Clampfit10.6. Action potential amplitude was calculated from threshold to peak. Action potential characteristics and passive cell membrane properties (cell capacitance, input resistance and resting membrane potential) were compared between genotypes using unpaired two-tailed Student’s t-test or Mann-Whitney test as specified in the text (*p* < 0.05). Input-output curve with firing frequency between heterozygous and wild-type mice was compared using two-way ANOVA (genotype and injected current as factors) with repeated measures (*p* < 0.05). All statistical tests were performed in Prism 8 (Graph Pad Software Inc.). All data sets were analyzed with D’Agostino Pearson normality test before carrying out parametric statistical tests, data sets that did not follow a normal distribution was compared using nonparametric Mann-Whitney test. Representative neuronal firing traces were plotted using MATLAB^5^ (MathWorks), all other data figures were plotted using Delta Graph 7.1 (RedRock Software).

## Results

### 1. Generation of Scn1a K1270T mutant mice

The GEFS+ associated K1270T mutation is located in the second transmembrane region of the third homology domain of the NaV1.1 sodium channel (Figure 1A, asterisk). The equivalent amino acid in mouse Nav1.1 is at K1259 in Nav1.1 (Figure 1B). To generate a knock-in mouse model, a 20 bp short guide RNA (sgRNA72, Figure 1B) was used (see Methods) to target the Cas9 cut site immediately upstream of the KT mutation in exon 19 (c.4227A) (Figure 1B). The repair template introduced a two base-pair mutation that converts a lysine (K: AAA) residue to threonine (T: ACC). For initial screening and routine genotyping, a silent mutation that results in an EcoRV restriction site was also included in the repair template (Figure 1B). DNA sequence modified by homology dependent repair is tracked by incorporation of silent mutations introduced via the repair template (Figure 1B). Following CRISPR mediated gene editing, *Scn1a*^*KT/+*^ allele carries K->T mutation, an EcoRV restriction enzyme cut site and two additional silent mutations to prevent Cas9 activity on the repaired allele. To remain consistent with the nomenclature in human subjects (Abou-Khalil et al.,2001), the mutant allele is referred to as *Scn1a*^*KT*^ (K1270T) (Figure 1B). Using standard protocols, guide RNAs and repair template were injected into mouse embryos that were transplanted into pseudo-pregnant CD1 females (Figure 1C).

**Figure 1.**
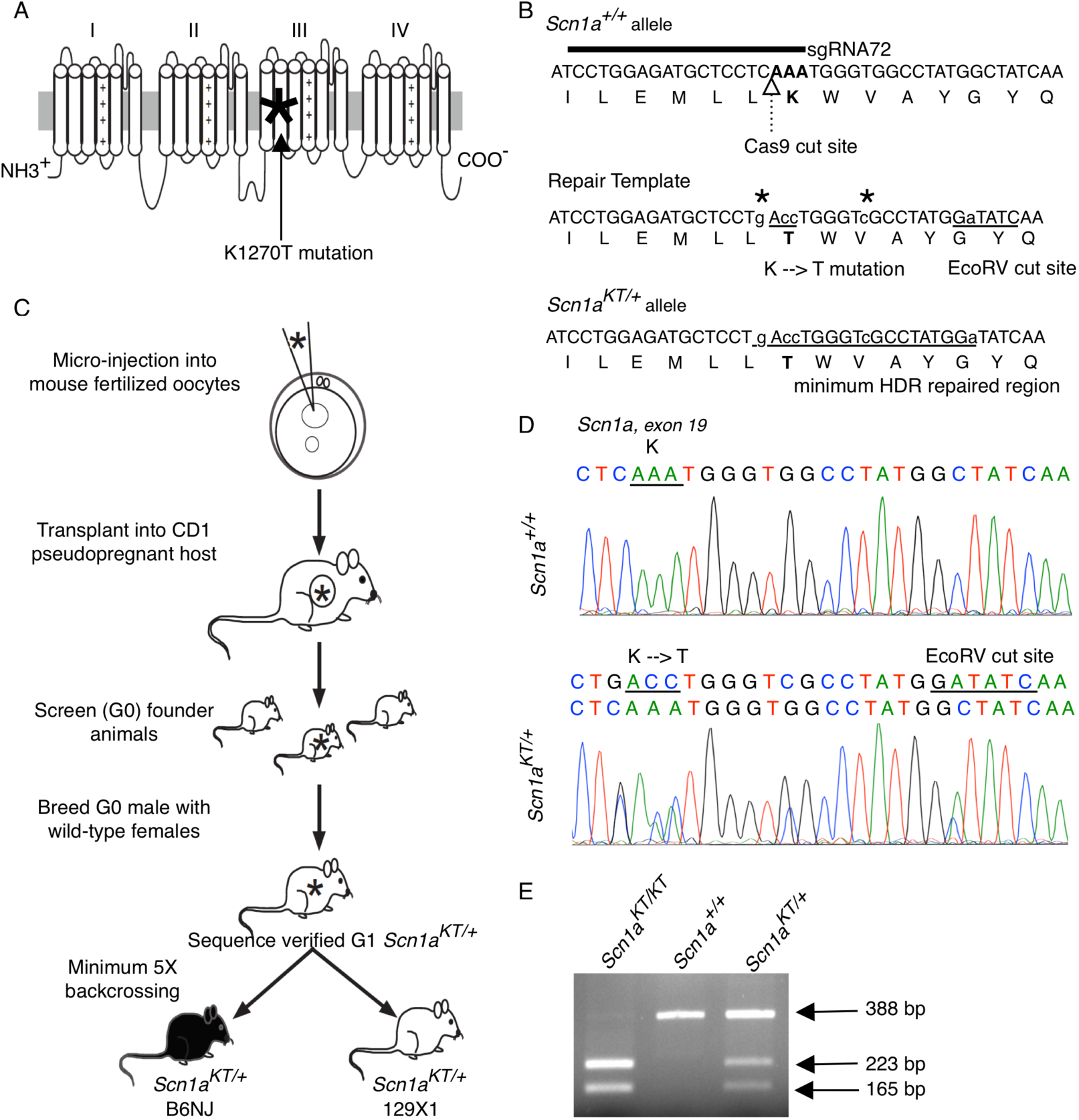
Generation of *Scn1a*^*KT/+*^ mouse using CRISPR/Cas9 technology. (A) GEFS+ causing K1270T mutation (asterisk) is located in S2 transmembrane segment of domain III of the alpha subunit of Nav1.1 ion channel encoded by the human *SCN1A* gene. (B) Location of the guide RNA relative to the Cas9 cut site and the locus of the K1259T mutation in the mouse *Scn1a* gene and the Nav1.1 protein sequence. Repair template sequence with the base pair changes introducing the K-T mutation and the EcoRV cut site. All edited nucleotides are shown in lowercase letters and the homology-dependent repair region is represented by underlined letters. Two additional silent mutations (asterisks) were added to prevent re-cutting by Cas9 following HDR. (C) Outline of the steps followed to generate transgenic K1270T mouse colonies in B6NJ and 129×1 genetic backgrounds via CRISPR/Cas9 gene editing. (D) DNA sequence comparison between a wild-type (*Scn1a*^*+/+*^) and a heterozygous (*Scn1a*^*KT/+*^) mouse showing missense K-T mutation and another silent mutation that results in an EcoRV cut site. (E) A representative agarose gel shows PCR amplified DNA bands digested with EcoRV which distinguishes between mice homozygous for the mutant allele, *Scn1a*^*KT/KT*^ (223 bp and 165 bp), *Scn1a*^*+/+*^ wild-type mice homozygous for the wild-type allele (388bp) and heterozygous *Scn1a*^*KT/+*^ mice carrying one copy each of wild-type (388 bp) and mutant (223 and 165 bp) alleles.

Previous studies have shown that the genetic background can influence seizure sensitivity in mice (Miller et al., 2013; Mistry et al., 2014; Ogiwara et al., 2007; Rubinstein et al., 2015b; Yu et al., 2006). To investigate whether strain background effects could be observed in *Scn1a*^*KT*^ mice, mutant mice were generated in two different strain backgrounds previously used to model *SCN1A* epilepsy disorders - C57BL/6NJ (or B6NJ) and 129×1/SvJ (or 129×1) by backcrossing for at least 5 generations (Figure 1C). Founder (G0) animals were screened to identify potential breeders that contained the mutant allele. Transmission of the mutant allele was verified by PCR amplification and sequencing to confirm the presence of both the K1270T mutation and EcoRV site at the appropriate locations in G1 and animals in subsequent generations (Figure 1D). Routine genotyping of animals took advantage of the EcoRV site to distinguish between wild-type, heterozygous and homozygous mutant animals (Figure 1E). Restriction digestion of the 388 bp PCR product from the mutant allele by EcoRV produced 223 bp and 165 bp products, while the wild-type allele 388 bp product is not digested.

### 2. *Scn1a*^*KT/KT*^ mice have reduced lifespan

Some *SCN1A* mutations that lead to either Dravet Syndrome (DS) or GEFS+ epilepsy have been correlated with premature death in mouse models (Yu et al., 2006; Ogiwara et al., 2007; Martin et al., 2010). To determine if the *Scn1a* K1270T mutation affected longevity, a life-span assay was conducted in both B6NJ and 129×1 strain backgrounds. Homozygous mutants (*Scn1a*^*KT/KT*^) in both backgrounds died prematurely between P17-P25 with average lifespan of 19.9 ± 0.5d and 20.3 ± 1.2d for 129×1 and B6NJ strains respectively (Figure 2A and 2B). Homozygous mice have a significantly shorter life-span than heterozygous and wild-type littermates in their respective background (Mantel Cox test, *df 2*, 129×1 strain: Chi square = 84.3, B6NJ: Chi square = 24.9, *p* < 0.0001) There was no difference in the lifespan of heterozygous (*Scn1a*^*KT/+*^) and wild-type (*Scn1a*^*+/+*^) littermates in the same background observed up to 6 months (Figure 2A and 2B).

**Figure 2.**
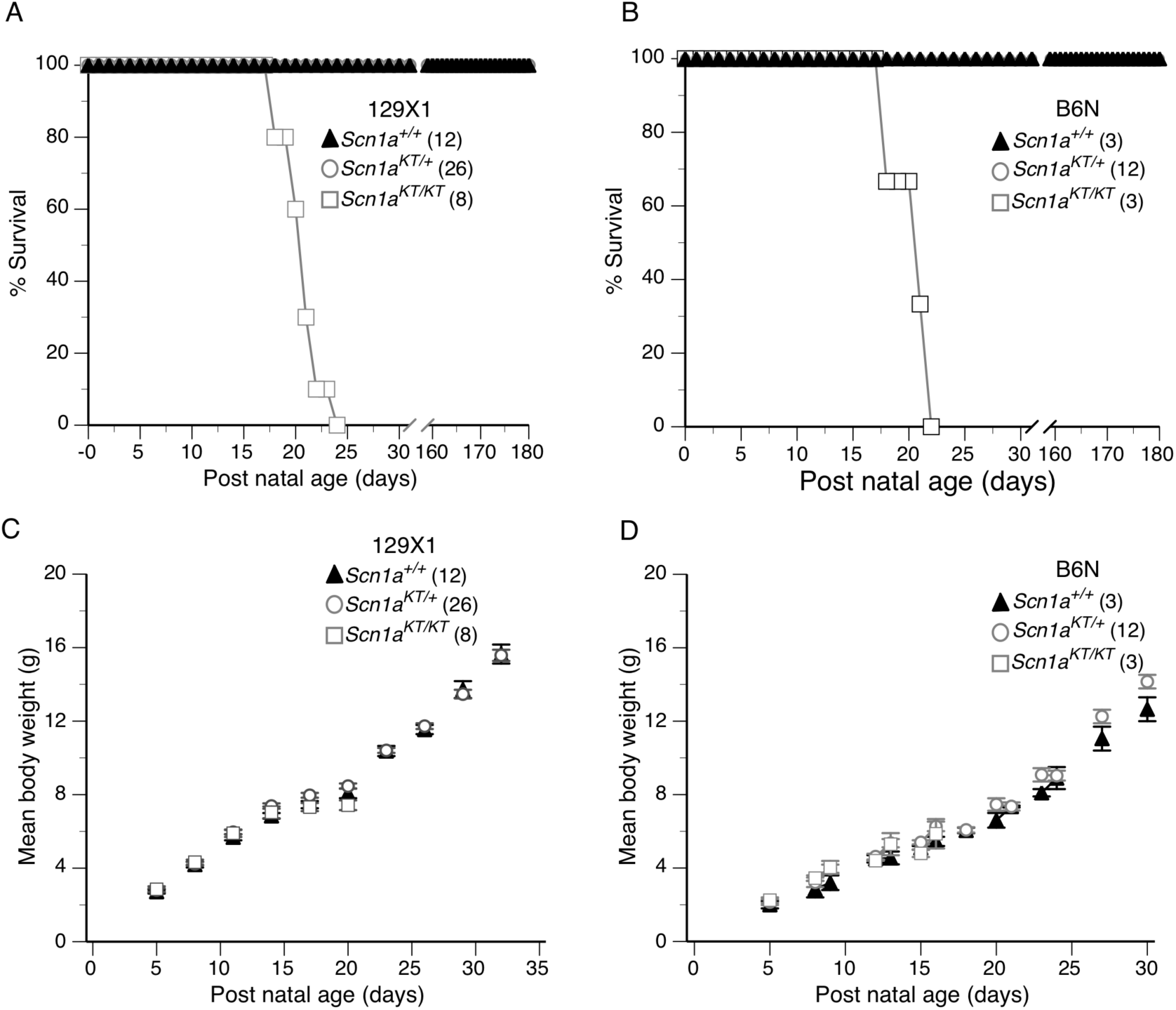
Homozygous (*Scn1a*^*KT/KT*^) mice have a shortened lifespan in both genetic backgrounds. (A,B) Survivorship plots of wild-type (*Scn1a*^*+/+*^) and mutant mice over a period of one month in 129×1 and B6NJ strains respectively. Homozygous (*Scn1a*^*KT/KT*^) mice in either background had a reduced mean lifespan (129×1 =19.9 days; B6NJ = 20.3 days) compared to heterozygous (*Scn1a*^*KT/+*^) and wild-type (*Scn1a*^*+/+*^) littermates assayed in parallel. (C,D) Mean body weight of mutant mice (*Scn1a*^*KT/+*^ and *Scn1a*^*KT/KT*^) was not different from wild-type (*Scn1a*^*+/+*^) littermates before the age of P20 (Mann-Whitney test, p < 0.05). Numbers in parantheses represent number of animals in each group. Data are presented as mean + S.E.M.

Spontaneous seizures were observed in some of the homozygous mice during daily monitoring and weighing of animals during the lifespan assay (*s*upplementary video V1). To determine if spontaneous seizure activity is correlated with death in the mutant mice, continuous video monitoring was conducted on three litters of 129×1 mice, between ages P18-P25, which included seven homozygous *Scn1a*^*KT/KT*^ pups and their littermates. All seven homozygous pups examined exhibited multiple spontaneous seizures with excessive jumping, limb jerking, falling on side, hind leg extension characteristics of classic tonic-clonic seizures (i.e., Racine score 5) *s*upplementary video V1). All seven homozygous pups died following severe spontaneous seizures. These results are similar to a previous study showing spontaneous seizures in homozygous *Scn1a*^*+/−*^ DS mice prior to death (Cheah et al., 2012).

Mean body weight of *Scn1a*^*KT/KT*^ homozygous mice in the 129×1 and B6NJ strain background increased at a similar rate to *Scn1a*^*KT/+*^ and *Scn1a*^*+/+*^ littermates between P5 and P16 (Figure 2C and 2D; Mann-Whitney test, *p > 0.05*). At P20, mean body weight of 129×1 *Scn1a*^*KT/KT*^ was significantly lower than heterozygous *Scn1a*^*KT/+*^ (Mann-Whitney test, *p* = 0.019) but not different from wild-type *Scn1a*^*+/+*^ mice (Mann-Whitney test, *p* = 0.147). Between P17-P21, homozygous mutant mice started dying in both 129×1 and B6NJ strain. Since there were less than three data points available to calculate mean body weight, no further statistical comparisons were performed on homozygous *Scn1a*^*KT/KT*^ mice on either strain. There was no difference in the mean body weight of heterozygous *Scn1a*^*KT/+*^ and wild-type *Scn1a*^*+/+*^ animals monitored up to P30-P32 either in 129×1 background (Figure 2C, two-way ANOVA, interaction *F* _(9,480)_ = 0.37, *p >* 0.05, followed by Sidak’s *post hoc* comparison) or B6N background (Figure 2D, two-way ANOVA, interaction *F* _(13,480)_ = 0.06, *p >* 0.05, followed by Sidak’s *post hoc* comparison). In heterozygous animals, the K1270T mutation does not affect rate of weight gain in the first month or life span up to 6 months in either strain. These data demonstrate that in homozygous condition, the K1270T mutation causes premature death in both 129×1 and B6NJ enriched backgrounds. Frequent spontaneous seizures could be the cause of death in homozygous mice as observed with pups in 129×1 background that were continuously video monitored.

### 3. *Scn1a*^KT/+^ mice exhibit heat-induced seizures

Detailed pedigree analysis of families with inherited GEFS+ has shown that individuals heterozygous for *SCN1A* mutations can exhibit a variety of seizure phenotypes with different clinical severity (Abou-Khalil et al., 2001; Scheffer, 1997; Zhang et al., 2017). However, febrile seizures are the most common feature of GEFS+ patients, including patients heterozygous for the K1270T mutation. To determine if the heterozygous mice phenocopy the heat-induced seizures in human patients, individual mice (P30-40) were fitted with a rectal temperature probe before being placed into a preheated custom-built chamber (Figure 3A, top panel). The chamber tempertaure was gradually increased by 5 °C every 10 min as illustrated (Figure 3A, bottom panel). The mean rate of rise in body temperature in response to heating of the chamber was similar between control *Scn1a*^*+/+*^ and heterozygous *Scn1a*^*KT/+*^ mice in both strain backgrounds (Figure 3B and 3C, two-tailed Student’s t-test, 129×1-*p* = 0.783; B6NJ-*p =* 0.318), indicating no overt changes in thermoregulation in *Scn1a*^*KT/+*^ mice.

**Figure 3.**
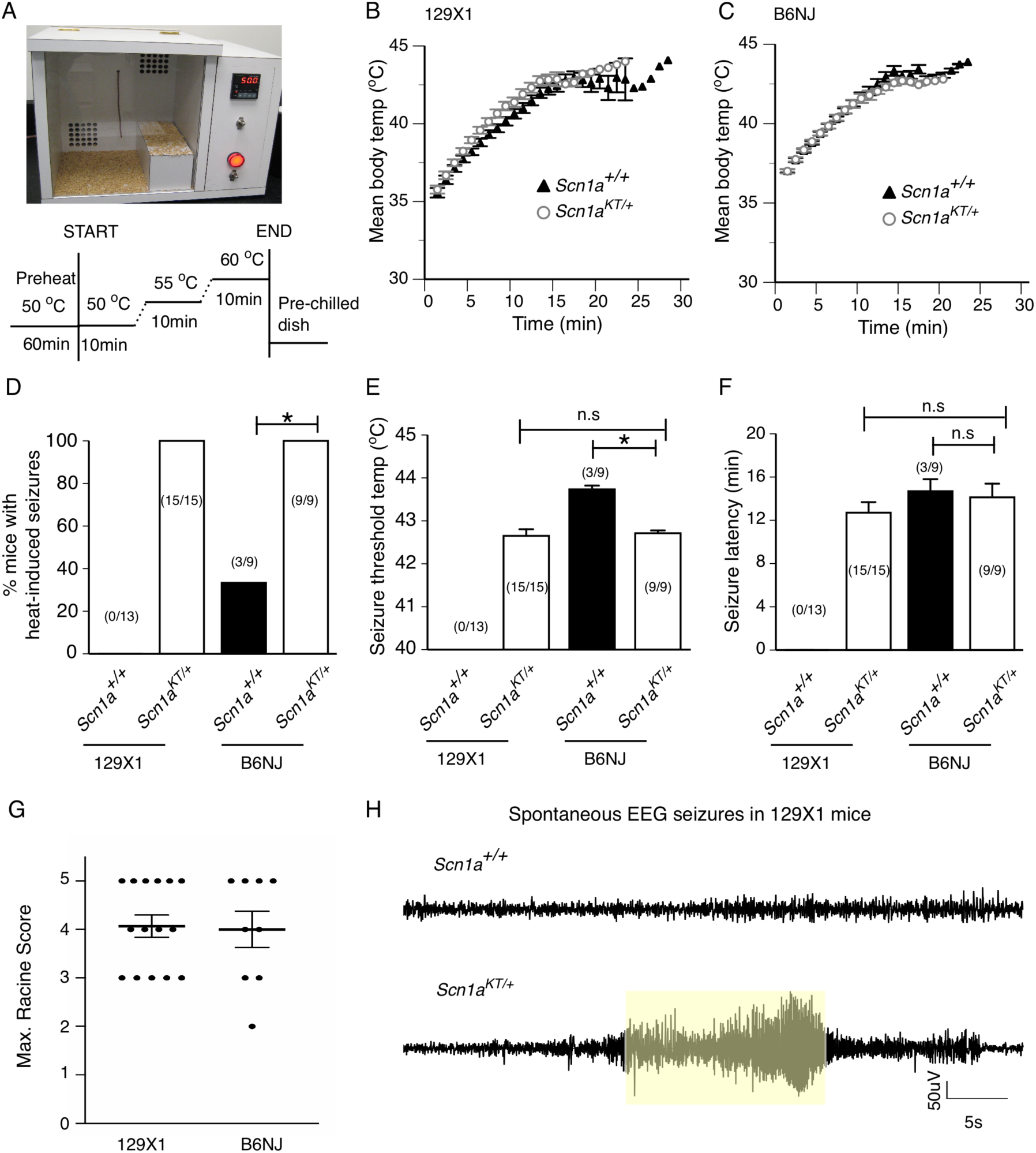
Heterozygous mice (*Scn1a*^*KT/+*^) exhibit heat induced seizures and spontaneous seizures. (A) Custom-built mouse heating chamber and schematic of the protocol used for heat-induced seizures. (B, C) Mean body temperature changes over time in wild-type (*Scn1a*^*+/+*^) and heterozygous mice (*Scn1a*^*KT/+*^) in 129×1 and B6NJ strains respectively. (D) Percentage of mice exhibiting heat-induced seizures in both strains. (E) seizure threshold temperature and (F) latency to heat-induced seizures between wild-type (*Scn1a*^*+/+*^) and heterozygous (*Scn1a*^*KT/+*^) mice in both strains. (G) Maximum Racine scores of heat-induced seizures exhibited by heterozygous (*Scn1a*^*KT/+*^) mice in both strains. Data are presented as mean + S.E.M. (H) Representative electrographic seizure traces from a 4-month old wild-type (*Scn1a*^*+/+*^) and heterozygous (*Scn1a*^*KT/+*^) mice on 129×1 strain. A recording trace from wild-types shows normal EEG pattern whereas a trace from a heterozygous mouse shows EEG pattern (shaded region) during a spontaneous seizure episode.

In the 129×1-enriched strain, no wild-type *Scn1a*^*+/+*^ mouse exhibited heat-induced seizures. In contrast, under identical conditions, all heterozygous *Scn1a*^*KT/+*^ mice had heat-induced seizures (Figure 3D). In the B6NJ background, a small percentage (33%) of the wild-type *Scn1a*^*+/+*^ mice exhibited heat-induced seizures while all heterozygous *Scn1a*^*KT/+*^ mice had heat-induced seizures (Figure 3D, supplementary video V2). The incidence of heat-induced seizures was significantly higher in heterozygous mice compared to wild-type mice in both backgrounds (Fisher’s exact test, 129×1 *p* < 0.0001; B6NJ *p* = 0.009). There was no difference between incidence of heat-induced seizures between heterozygous *Scn1a*^*KT/+*^ mice in either 129×1 or B6NJ background (Figure 3D; Fisher’s exact test, *p* > 0.999).

Average seizure threshold temperature was similar in the heterozygous *Scn1a*^*KT/+*^mice in both backgrounds: 129×1 - 42.6 ± 0.20 °C, B6NJ - 42.7 ± 0.06 °C (Figure 3E; two-tailed unpaired Student’s t-test, *p* = 0.782). In the small percentage of B6NJ wild-type mice that did have seizures (3 out of 9 mice), the mean seizure threshold temperature was significantly higher than the heterozygous mice: *Scn1a*^*+/+*^ - 43.7 ± 0.08 °C vs *Scn1a*^*KT/+*^ - 42.7 ± 0.06 °C (Figure 3E, two-tailed unpaired Student’s t-test, *p* < 0.0001). There was no difference in the latency to seizure onset between heterozygous mice on either strains: 129×1 - 12.7 ± 0.9min, B6NJ - 14.1 ± 1.2 min (Figure 3F; Mann-Whitney test, *p* = 0.290). Similarly, latency to seizure was not different between wild-type (*Scn1a*^*+/+*^ - 14.7 ± 1.1 min) and heterozygous mice (*Scn1a*^*KT/+*^ −14.1 ± 1.2 min) on B6NJ background (Figure 3E, two-tailed unpaired Student’s t-test, *p* = 0.813). Also, maximal seizure severity as measured on modified Racine scale between heterozygus *Scn1a*^*KT/+*^ mice in the two backgrounds was not different (Figure 3G; Mann-Whitney test, *p* > 0.9999). The heterozygous mice in both strains had a mean Racine score of 4 (Figure 3G). Mice would typically begin to exhibit heat-induced seizures with facial spasms or head-nodding (Racine score 1-2) and gradually progress to forelimb clonus, falling on sides, hindlimb extension and/or uncontrolled jumping (Racine score 5).

In a small number of trials, heterozygous mice that seized at ~42 °C were left in the chamber until their body tempertaure reached 43 °C - 44 °C, to mimic the average seizure threshold of B6NJ wild-type *Scn1a*^*+/+*^ animals. This resulted in a second, more severe seizure, characterized by uncontrolled jumping, all limbs jerking (Racine score 5) followed by immediate death of the mice, indicating mutant mice were more sensitive to heat-induced seizures. Taken together, our data demonstrate that all heterozygous mutant mice exhibit heat-induced seizures with similar frequency, seizure threshold, latency and seizure severity in a strain-independent manner.

### 4. Spontaneous seizures in heterozygous Scn1a^KT/+^ mice

Some GEFS+ patients with *SCN1A* K1270T mutation have afebrile seizures in addition to febrile seizures (Abou-Khalil et al., 2001). Therefore the *Scn1a*^*KT/+*^ mice were screened for occurrence of spontaneous seizures via continuous EEG monitoring over a seven-day period. EEG monitoring was performed on adult mice on the 129×1-enriched background between the age of 3-4 months. One out of five heterozygous (*Scn1a*^*KT/+*^) animals exhibited spontaneous seizures whereas none of the wild-type littermate (*Scn1a*^*+/+*^) animals examined in parallel displayed spontaneous seizures. Representative EEG recordings from the heterozygous mouse that had spontaneous seizures reveals a classic electrographic seizure waveform with high amplitude and high frequency polyspike discharges (Figure 3H). This animal experienced an average of 18.1 ± 5.3 seizures per day with a mean seizure duration of 38.9 ± 6.2s. This suggests that the heterozygous mutation results in spontaneous seizures in mice, consistent with the patient phenotype where individuals with K1270T or other GEFS+ mutations exhibit afebrile, tonic-clonic generalized seizures in low incidence (Abou-Khalil et al., 2001; Zhang et al., 2017).

### 5. Impaired excitability of parvalbumin-positive interneurons

Mutations in *Scn1a* are associated with reduced excitability of PV-expressing inhibitory interneurons in mouse models of Dravet Syndrome (Cheah et al., 2012; Ogiwara et al., 2007; Rubinstein et al., 2015a; Tai et al., 2014; Yu et al., 2006) and GEFS+ (R1648H) disorders (Hedrich et al., 2014; Martin et al., 2010). Reduced excitability of inhibitory neurons had been implicated in a *Drosophila* model of K1270T mutation as well (Sun et al., 2012). To determine the effect of K1270T mutation on PV interneurons, *Scn1a*^*KT/+*^ mice were crossed with PV-Cre mice and Ai14-tdTomato reporter on a C57BL/6J background (i.e., to generate *Scn1a*^*KT/+*^;PV-Cre;tdTomato mice). Genetic labeling of PV neurons with tdTomato (magenta) was confirmed by co-immunostaining brain slices with anti-PV antibody (green) and nuclei marker DAPI (blue) (Figure 4A, white arrows). Subsequently, we obtained current-clamp recordings from tdTomato+ neurons in CA1 of wild-type and *Scn1a*^*KT/+*^ mice at P30-40.

**Figure 4.**
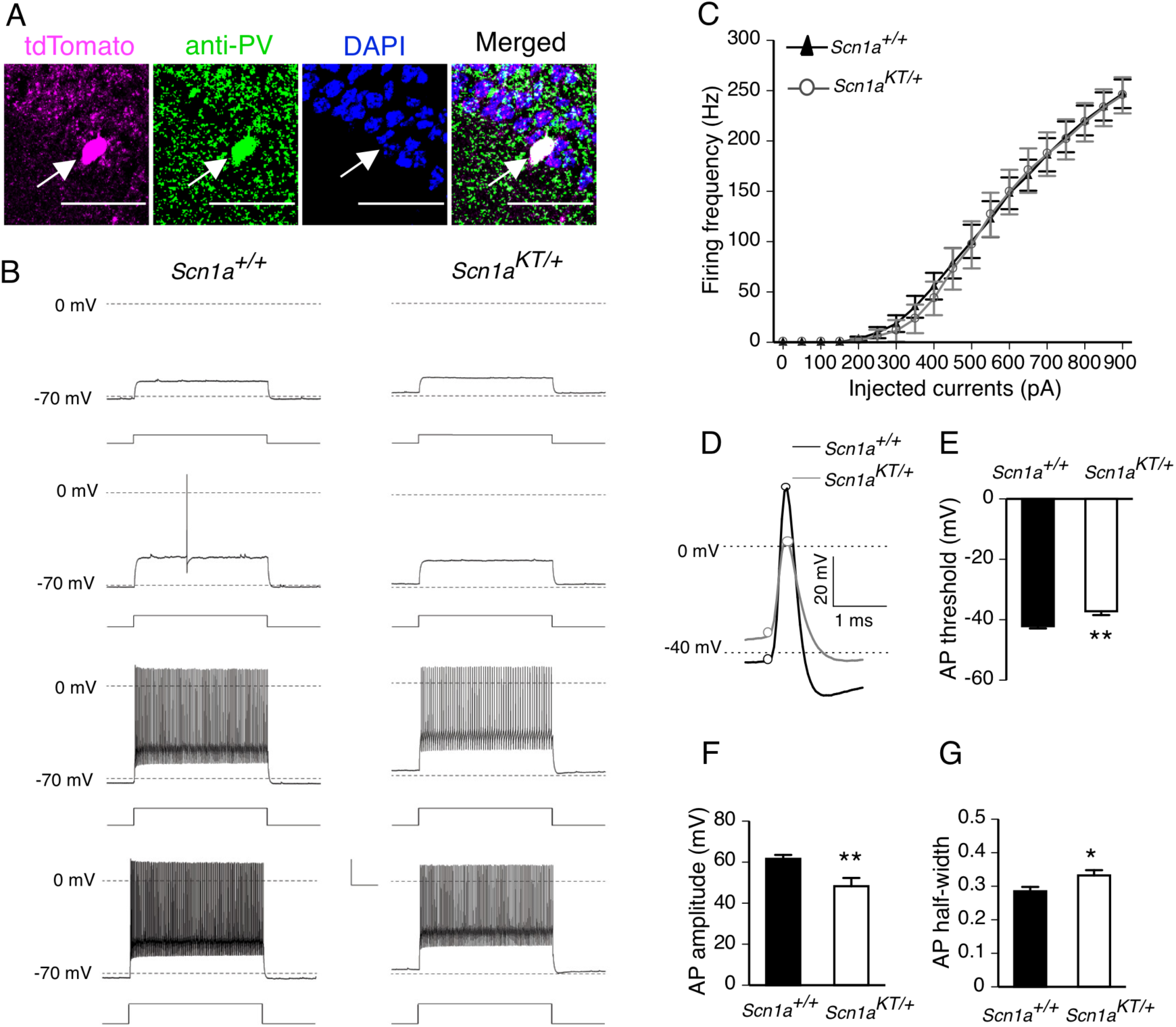
Reduced excitability of PV neurons in *Scn1a*^*KT/+*^ mice. (A) A representative PV interneuron (arrow) which was genetically labeled with tdTomato (magenta) in *Scn1a*^*KT/+*^;*PV-Cre;Ai14-tdTomato* mice and co-immunostained with anti-PV antibody (green) and DAPI (blue). Right most panel shows the merged image. Scale bar is 50 µM. (B) Representative traces of PV interneurons from wild-type (*Scn1a*^*+/+*^) and heterozygous (*Scn1a*^*KT/+*^) littermates at different current injection steps. (C) Input-output curves showing AP firing frequency in PV interneurons against current injection steps between 0 pA and 900 pA, data point for 50 pA step increment is shown here. No difference in PV firing frequency between *Scn1a*^*KT/+*^ (grey curve, open circles) and wild-type *Scn1a*^*+/+*^ mice (black curve, triangles) was seen (two-way ANOVA with repeated measures, *p* > 0.05). (D) Expanded representative traces of first AP fired from a PV interneuron recorded from a *Scn1a*^*+/+*^ and *Scn1a*^*KT/+*^ mouse. (E) AP threshold is more depolarized in *Scn1a*^*KT/+*^ mice compared to wild-type *Scn1a*^*+/+*^ mice (two-tailed unpaired t-test, *p* = 0.001). (F) AP amplitude is also reduced in *Scn1a*^*KT/+*^ mice compared to *Scn1a*^*+/+*^ littermates (two-tailed unpaired t-test, *p* = 0.030). (F) AP half-width is increased in *Scn1a*^*KT/+*^ mice (two-tailed unpaired t-test, *p* = 0.031). Data shown are average of 14 and 9 cells from *Scn1a*^*+/+*^ and *Scn1a*^*KT/+*^ littermates respectively recorded from at least 3 different mice. Error bars are S.E.M.

Characteristic fast spiking firing patterns were observed in td-tomato+ neurons in both wild-type *Scn1a*^*+/+*^ and heterozygous *Scn1a*^*KT/+*^ mice (Figure 4B), suggesting fast spiking properties remain intact in the mutants. Input-output curve showing firing frequency of PV interneurons was not significantly different between heterozygous *Scn1a*^*KT/+*^ (gray, open circles) and wild-type *Scn1a*^*+/+*^ control mice (black, triangles) (Figure 4C, two-way ANOVA with repeated measures, interaction: *F*_(90, 1890)_ = 0.1346, *p* > 0.9999). Single action potential properties of PV interneurons were also compared and interestingly, there were differences in action potential characteristics between heterozygous *Scn1a*^*KT/+*^ and wild-type *Scn1a*^*+/+*^ mice (Table 1, Figure 4D-4G). Although both wild-type and heterozygous PV interneurons fired typical brief action potentials with a rapid afterhyperpolarization (Figure 4D), action potential threshold in heterozygous *Scn1a*^*KT/+*^ was more depolarized potential than wild-type *Scn1a*^*+/+*^ mice (Table 1, Figure 4E, two-tailed unpaired Student’s t-test, *p* = 0.001). Also, action potential amplitude in heterozygous *Scn1a*^*KT/+*^ was reduced compared to wild-type *Scn1a*^*+/+*^ mice (Table 1, Figure 4D, Figure 4F, two-tailed unpaired Student’s t-test, *p* = 0.003). Action potential half-width was broader in *Scn1a*^*KT/+*^ different compared to control *Scn1a*^*+/+*^ mice (Figure 4G). All other intrinsic membrane properties including RMP was not significantly different between *Scn1a*^*+/+*^ and *Scn1a*^*KT/+*^ mice (Table 1). These results suggest that K1270T GEFS+ mutation leads to significant impairment in action potential firing of PV neurons without altering input-output firing properties.

**Table 1.** Electrophysiological properties of PV interneurons

| Properties | *Scn1a*<sup>+/+</sup> | *Scn1a*<sup>KT/+</sup> | *p* value |
| --- | --- | --- | --- |
| cells # (animals #) | 14 (5) | 9 (3) |  |
| Capacitance (pF) | 45.58 ± 9.16 | 60.0 ± 18.69 | 0.402<sup>a</sup> |
| Input resistance (MΩ) | 111.79 ± 21.86 | 74.83 ± 8.94 | 0.477<sup>a</sup> |
| Resting potential (mV) | -65.14 ± 0.61 | -63.88 ± 0.82 | 0.227 |
| AP threshold (mV) | -42.14 ± 0.77 | -37.44 ± 1.17** | 0.002 |
| AP amplitude (mV) | 61.60 ± 1.91 | 46.68 ± 3.93** | 0.003 |
| AP half-width (ms) | 0.285 ± 0.01 | 0.332 ± 0.01* | 0.031 |

Statistical comparisons were made with unpaired<sup>a</sup> Mann Whitney test or Student's *t*-test with \* = $p < 0.05$ .

### 6. Excitability of CA1 pyramidal neurons is unaltered

Although hyperexcitability in SCN1A epilepsy mouse models has been largely associated with impaired firing in inhibitory neurons (Cheah et al., 2012; Dutton et al., 2013; Hedrich et al., 2014; Martin et al., 2010; Ogiwara et al., 2007; Rubinstein et al., 2015a; Tai et al., 2014; Yu et al., 2006), several studies suggests that excitatory neuronal firing could also be altered as seen in a human iPSC *SCN1A* model (Liu et al., 2013) and *SCN2A* mouse model (Ogiwara et al., 2018). Action potential firing was recorded in current-clamp mode from excitatory CA1 pyramidal neurons from adult (P28-P40) mouse brain slices. Representative traces from wild-type *Scn1a*^*+/+*^ and heterozygous *Scn1a*^*KT/+*^ mice show regular spiking firing pattern with characteristic AP-doublet seen at 100 pA step and beyond (Figure 5A). Input-output curve shows that excitatory firing frequency is not significantly different in heterozygous *Scn1a*^*KT/+*^ mice (grey curve, open circles) compared to wild-type *Scn1a*^*+/+*^ littermates (black curve, triangles) (Figure 5B, two-way ANOVA with repeated measures, interaction: *F*_(28, 756)_ = 0.6038, *p* = 0.948). Individual action potential traces appear indistinguishable between *Scn1a*^*+/+*^ and heterozygous *Scn1a*^*KT/+*^ mice (Figure 5C). Action potential characteristics such as threshold, amplitude or half-width was not significantly different between heterozygous *Scn1a*^*KT/+*^ and *Scn1a*^*+/+*^ control mice (Table 2, Figure 5D-5F, two-tailed unpaired Student’s t-test, *p* > 0.05). Cell membrane properties such as cell capacitance, input resistance or resting membrane potential (RMP) was also not different between genotypes (Table 2, two-tailed unpaired Student’s t-test, *p* > 0.05). These results suggest that there is no change in neuronal firing in CA1 excitatory neurons in heterozygous *Scn1a*^*KT/+*^ mice consistent with other *Scn1a* mouse models (Cheah et al., 2012; Dutton et al., 2013; Hedrich et al., 2014; Martin et al., 2010; Ogiwara et al., 2007; Rubinstein et al., 2015a; Tai et al., 2014; Yu et al., 2006).

**Table 2.** Electrophysiological properties of excitatory CA1 pyramidal neurons

| Properties | *Scn1a*<sup>+/+</sup> | *Scn1a*<sup>KT/+</sup> | *p* value |
| --- | --- | --- | --- |
| cells # (animals #) | 12 (8) | 17 (9) |  |
| Capacitance (pF) | 27.71 ± 1.93 | 31.77 ± 3.23 | 0.340 |
| Input resistance (MΩ) | 182.04 ± 20.59 | 187.62 ± 13.58 | 0.815 |
| Resting potential (mV) | -65.16 ± 1.01 | -65.09 ± 0.69 | 0.947 |
| AP threshold (mV) | -45.29 ± 1.27 | -46.12 ± 1.51 | 0.695 |
| AP amplitude (mV) | 79.57 ± 3.26 | 72.45 ± 3.91 | 0.199 |
| AP half width (ms) | 2.16 ± 0.14 | 2.42 ± 0.15 | 0.238 |

Statistical comparisons were made with two-tailed unpaired t-test with \* = $p < 0.05$ .

**Figure 5.**
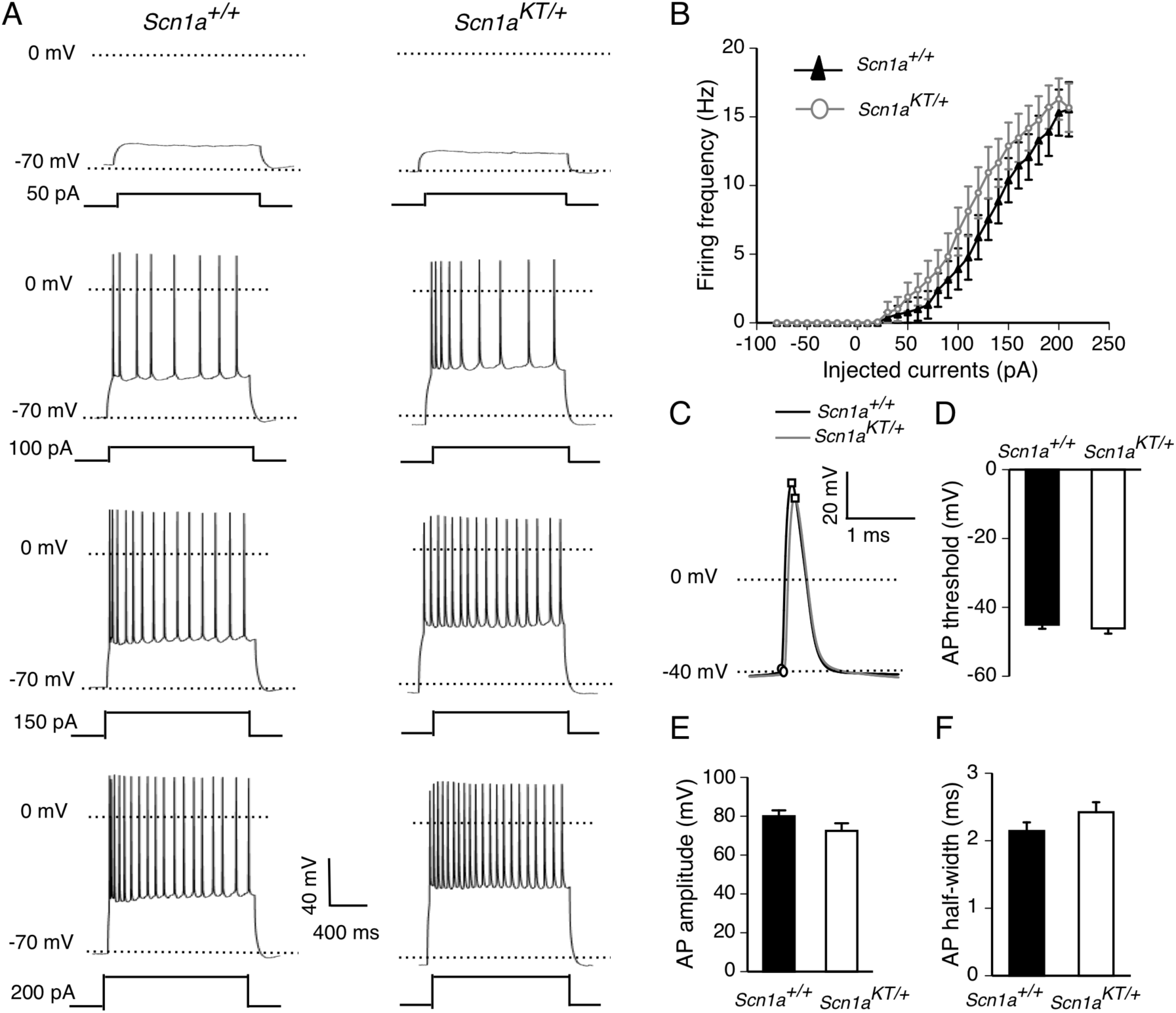
Firing property of CA1 excitatory neurons in *Scn1a*^*KT/+*^ mice remains unaltered. (A) Representative traces of action potential firing from CA1 excitatory cells in wild-type (*Scn1a*^*+/+*^) and heterozygous (*Scn1a*^*KT/+*^) mice in response to increasing current injection steps. (B) No difference in firing frequency found between heterozygous *Scn1a*^*KT/+*^ mice compared and *Scn1a*^*+/+*^ littermates (two-way ANOVA, *p* > 0.05). (C) Expanded single AP traces from *Scn1a*^*+/+*^ and *Scn1a*^*KT/+*^ mice (D-F) No change observed in RMP or AP threshold, AP amplitude, AP half-width between heterozygous *Scn1a*^*KT/+*^ and *Scn1a*^*+/+*^ mice (two-tailed unpaired t-test, *p* > 0.05). Data shown is average of 12 and 17 cells from *Scn1a*^*+/+*^ and *Scn1a*^*KT/+*^ littermates respectively recorded from at least 7 different mice. Error bars are S.E.M.

## Discussion

### Seizure behaviors and genetic background

To our knowledge, this is the first study to successfully use CRISPR/Cas9-based gene-editing technique to generate a mouse model of human GEFS+ epilepsy. We report that approximately 20% of heterozygous *Scn1a*^*KT/+*^ mice on a 129×1 genetic background show spontaneous seizures at ambient room temperature. Such sporadic incidence of spontaneous seizures has been previously reported in other *Scn1a* mouse models: 3 out of 7 mice in *Scn1a*^*+/−*^ DS model (Yu et al., 2006) and 2 out of 14 mice in *Scn1a*^*RH/+*^ GEFS+ mouse model (Martin et al., 2010). It is also consistent with the low incidence rate of afebrile seizures found in a larger population of human patients (Zhang et al., 2017). In contrast, all heterozygous mice in either 129×1 or B6NJ background exhibit heat-induced seizures at a seizure threshold temperature (42-43 °C) similar to R1648H *Scn1a*^*RH/+*^ mice on C57BL/6J background (Martin et al., 2010). This is consistent with the phenotypes in human patients in which the FS/FS+ is the predominant feature of GEFS+ disorder (Zhang et al., 2017).

Homozygous *Scn1a*^*KT/KT*^ mice display recurrent spontaneous seizures prior to death. Sudden unexpected death in epilepsy (SUDEP) is known to occur possibly due to lethal cardiac arrhythmias following severe seizures in *Scn1a* epilepsy mouse model (Auerbach et al., 2013; Kalume et al., 2013). Simultaneous EEG and electrocardiogram (ECG) monitoring on homozygous *Scn1a*^*KT/KT*^ mice could help us determine if cardiac dysfunction results in death following spontaneous seizures. It will also be interesting to examine if heat-induced seizures in juvenile homozygous mutant mice (evaluated prior to their death) is more dramatic to heterozygous mice in terms of reduced seizure threshold and seizure latency as seen in R1648H GEFS+ mouse model (Martin et al,.2010).

Previous studies have reported that mutant *Scn1a*^*+/−*^ mice on C57BL/6J background show a higher susceptibility to spontaneous seizures and reduced lifespan, in contrast to mutant mice on 129×1/SvJ background (Yu et al., 2006). Effects of lifespan and heat-induced seizures have been studied only on C57BL/6J background in R1648H GEFS+ mouse at N2 generation (Martin et al., 2010) and at N12 backcross generations (Sawyer et al., 2016). Mutant K1270T mice used in this study were backcrossed at least five times (N5) to their respective genetic backgrounds and no strain-dependent differences was observed in lifespan and heat-induced seizure susceptibility. It is possible that genetic modifiers may influence strain-dependent seizure phenotypes (Miller et al., 2013; Ogiwara et al., 2007; Yu et al., 2006) such as other ion channels (Bergren et al., 2009; Hawkins et al., 2011; Jorge et al., 2011) or transcription factors like hepatic leukemia factor (*Hlf*) (Hawkins and Kearney, 2016). It is also possible that strain dependent seizure susceptibility in K1270T mice may be seen at a later age, similar to increased susceptibility to chemoconvulsants such as kainic acid seen in older mice in other seizure models (McCord et al., 2008). Future studies will include monitoring seizure generation in mice of two genetic background at a prolonged time course.

### Effects of SCN1A mutations on neuronal firing

These results demonstrate selective impairment of evoked firing in hippocampal PV-expressing inhibitory neurons, while the firing properties of pyramidal excitatory neurons are unaltered. The mouse hippocampus contains a heterogenous population of inhibitory GABAergic interneurons, including PV, somatostatin (SST) and vasoactive intestinal peptide (VIP)-expressing neurons, all that play an important role in microcircuits (Pelkey et al., 2017; Soltesz and Losonczy, 2018). Dysfunction in one or more of these inhibitory neuron populations can disrupt both the feedback and feedforward inhibitory circuitry, resulting in epilepsy disorders (Khoshkhoo et al., 2017; Paz and Huguenard, 2015). NAV1.1 is predominantly localized in the soma and in the axon initiating segment of PV-positive neurons (Ogiwara et al., 2007) and to lesser extent in SST and VIP neurons (Goff and Goldberg, 2019; Li et al., 2014; Ogiwara et al., 2007; Tai et al., 2014). Reduced excitability of PV neurons has been implicated in hyperexcitable circuits in *Scn1a* mouse models of DS and GEFS+ (Dutton et al., 2013; Favero et al., 2018; Hedrich et al., 2014; Rubinstein et al., 2015a; Tai et al., 2014). Selective deletion of Nav1.1 in PV or SST-expressing neurons resulted in reduced neuronal excitability but led to phenotypically distinct behavioral disorders, suggesting distinct roles of interneurons subpopulations in mediating DS phenotypes (Rubinstein et al., 2015a). Recently, irregular spiking of VIP interneurons has been associated with DS phenotypes in an *Scn1a*^*+/−*^ mouse model (Goff and Goldberg, 2019), emphasizing the need to focus on multiple subtypes of interneuron population to more thoroughly understand how impaired neuronal firing contributes to seizure phenotypes in GEFS+ and other *Scn1a* mouse models.

In a R1648H GEFS+ mouse model, excitatory neuronal firing remained unchanged whereas bipolar inhibitory neuronal firing was reduced in mutant mice compared to controls (Martin et al., 2010). In the K1270T mouse model, there is no gross reduction in firing frequency of either excitatory pyramidal cells or PV GABAergic interneurons. However, PV interneuron excitability is altered in heterozygous *Scn1a*^*KT/+*^ mice as reflected in reduced AP amplitude and depolarized AP threshold. A recent study showed that decreased firing frequency in PV interneurons in DS *Scn1a*^*+/−*^ mutant mice in an early age window (P18-21) but no detectable change in firing frequency at a later age (P35-36) (Favero et al., 2018). Future studies that examine firing properties of subpopulations of GABAergic interneurons at developmentally critical age windows may help determine how the K1270T mutation alters the functional development of interneurons. In addition, in order to understand the alterations of sodium current properties in K1270T mutant mice, it might be informative to investigate if alterations in sodium channel function are restricted to distinct subtypes of inhibitory neurons. It will also be important to determine if there are exacerbated or additional defects in the evoked firing of PV and other GABAergic interneurons at high temperatures in the mutant mice.

### Comparison with the existing models of K1270T mutation

Elucidating the cellular machinery in taxonomically diverse models can help identify conserved pathway that leads to seizure phenotypes induced by the GEFS+ causing K1270T mutation. At the behavioral level, both K1270T knock-in flies and heterozygous *Scn1a*^*KT/+*^ mice exhibited heat-seizures and low incidence of spontaneous seizures (Sun et al., 2012). This is consistent with K1270T GEFS+ phenotype found in human patients (Abou-Khalil et al., 2001). In both fly and hiPSC models of K1270T, the mutation causes reduced firing in inhibitory neurons and increased firing in excitatory neurons in heterozygotes, although the deficits in fruit-flies occur at elevated temperatures and those in human iPSC-derived neurons at room temperature. In addition, in the K1270T mutant flies, reduced firing in inhibitory neurons is driven by a temperature dependent hyperpolarized shift in deactivation threshold of persistent sodium currents (Sun et al., 2012), while reduced firing in iPSC-derived inhibitory neurons results from reduction in sodium current density (Xie et al., 2019). In the current study of K1270T mouse, PV inhibitory firing frequency is not reduced however the AP threshold is shifted to a more depolarized potential and AP amplitude is significantly reduced. This suggests that the K1270T mutation makes PV interneurons less excitable. In contrast to the K1270T flies and the heterozygous hiPSC cell line, excitatory neuronal firing remains unaltered in heterozygous mice compared to wild-types. However, this is consistent with many other studies in mouse models of DS and GEFS+ that show unaltered excitability of pyramidal neurons (Cheah et al., 2012; Dutton et al., 2013; Martin et al., 2010; Ogiwara et al., 2007; Rubinstein et al., 2015a; Tai et al., 2014; Yu et al., 2006).

The ultimate goal of our study is to develop mechanism-based drug therapies using our mouse model that recapitulates seizure phenotypes in human patients. The micro-electrode array (MEA) system is emerging as a powerful tool to examine long term neural network activity in rodent brain slices or neuronal cultures in response to pharmacological drugs (Jensen, 2019; Liu et al., 2012; McConnell et al., 2012; Valdivia et al., 2014). Previous studies have shown that the MEA can be utilized for recording spontaneous neuronal activity from acute hippocampal cultures derived from R1648H mouse (Hedrich et al., 2014) and screen anti-convulsant drugs in rat brain slices (Fan et al., 2019). Our aim is to establish a high-throughput platform using MEA to screen potential anti-convulsant drug candidates that suppress aberrant network activity in the GEFS+ mouse model. Overall, mouse models of *SCN1A* epilepsy that are generated by the CRISPR/Cas9-based gene-editing methods advance our understanding on the cellular and circuit mechanism of *SCN1A* mutations and facilitate the development of anti-convulsant drugs.

## Acknowledgement

This research was supported by funds from NIH grants to D.O.D (NS083009) and to R.F.H (NS096012). We are thankful to Xiangmin Xu (UCI) for sharing *PV-Cre* mouse line (JAX stock# 017320) with us. We would like to thank Martin Smith and Connor Smith for constructing the mouse heat chamber box, Andrzej Foik for 3-D printing the brain slice incubation chamber, Andrew Salgado and Daniel Benavides for help with mouse genotyping, and Longwen Huang for help with MATLAB codes. We acknowledge the support of the Chao Family Comprehensive Cancer Center Transgenic Mouse Facility Shared Resource, supported by the National Cancer Institute of the National Institutes of Health under award number P30CA062203.

## Author contributions

S.S, A.D and D.K.O conceptualized the study; G.R.M and J.C.N designed the CRISPR strategy and generated the K1270T mouse line; A.D and L.Z performed the heat-induced seizure assay and immunostaining; A.D and A.P conducted EEG surgeries; A.D, B.Z, Y.X performed electrophysiological studies; A.D performed the lifespan assay, analyzed data, drafted the figures and wrote the manuscript. G.R.M, R.F.H, Y.X and D.O.D edited the manuscript.

